# Structural basis for functional properties of cytochrome *c* oxidase

**DOI:** 10.1101/2023.03.20.530986

**Authors:** Izumi Ishigami, Raymond G. Sierra, Zhen Su, Ariana Peck, Cong Wang, Frederic Poitevin, Stella Lisova, Brandon Hayes, Frank R. Moss, Sébastien Boutet, Robert E. Sublett, Chun Hong Yoon, Syun-Ru Yeh, Denis L. Rousseau

## Abstract

Cytochrome *c* oxidase (C*c*O) is an essential enzyme in mitochondrial and bacterial respiration. It catalyzes the four-electron reduction of molecular oxygen to water and harnesses the chemical energy to translocate four protons across biological membranes, thereby establishing the proton gradient required for ATP synthesis^1^. The full turnover of the C*c*O reaction involves an oxidative phase, in which the reduced enzyme (**R**) is oxidized by molecular oxygen to the metastable oxidized **O**_**H**_ state, and a reductive phase, in which **O**_**H**_ is reduced back to the **R** state. During each of the two phases, two protons are translocated across the membranes^2^. However, if **O**_**H**_ is allowed to relax to the resting oxidized state (**O**), a redox equivalent to **O**_**H**_, its subsequent reduction to **R** is incapable of driving proton translocation^2,3^. How the **O** state structurally differs from **O**_**H**_ remains an enigma in modern bioenergetics. Here, with resonance Raman spectroscopy and serial femtosecond X-ray crystallography (SFX)^4^, we show that the heme *a*_3_ iron and Cu_B_ in the active site of the **O** state, like those in the **O**_**H**_ state^5,6^, are coordinated by a hydroxide ion and a water molecule, respectively. However, Y244, a residue covalently linked to one of the three Cu_B_ ligands and critical for the oxygen reduction chemistry, is in the neutral protonated form, which distinguishes **O** from **O**_**H**_, where Y244 is in the deprotonated tyrosinate form. These structural characteristics of **O** provide new insights into the proton translocation mechanism of C*c*O.

## Main

Mammalian C*c*O is a large integral membrane protein comprised of 13 subunits. It contains four redox active centers, Cu_A_, heme *a*, and a heme *a*_3_/Cu_B_ binuclear center (BNC) (Fig. 1A). Molecular oxygen binds to the heme *a*_3_ iron in the BNC, where it is reduced to two water molecules by accepting four electrons from cytochrome *c* and four protons (the “substrate” protons) from the negative side (N-side) of the mitochondrial membrane (Fig. 1A). The energy derived from the oxygen reduction chemistry is used to drive the translocation of four protons (the “pumped” protons) from the N-side to the positive side (P-side) of the membrane^1,7^. Strong evidence suggests that the substrate protons are delivered to the BNC *via* the D and K-channel (see Extended Data Fig. 1), while the pumped protons are translocated *via* the D-channel^8-10^ or the H-channel^11^ through a putative proton loading site (PLS) located between the heme *a*_3_ propionates and a Mg^2+^ center^12-15^.

**Fig. 1.**
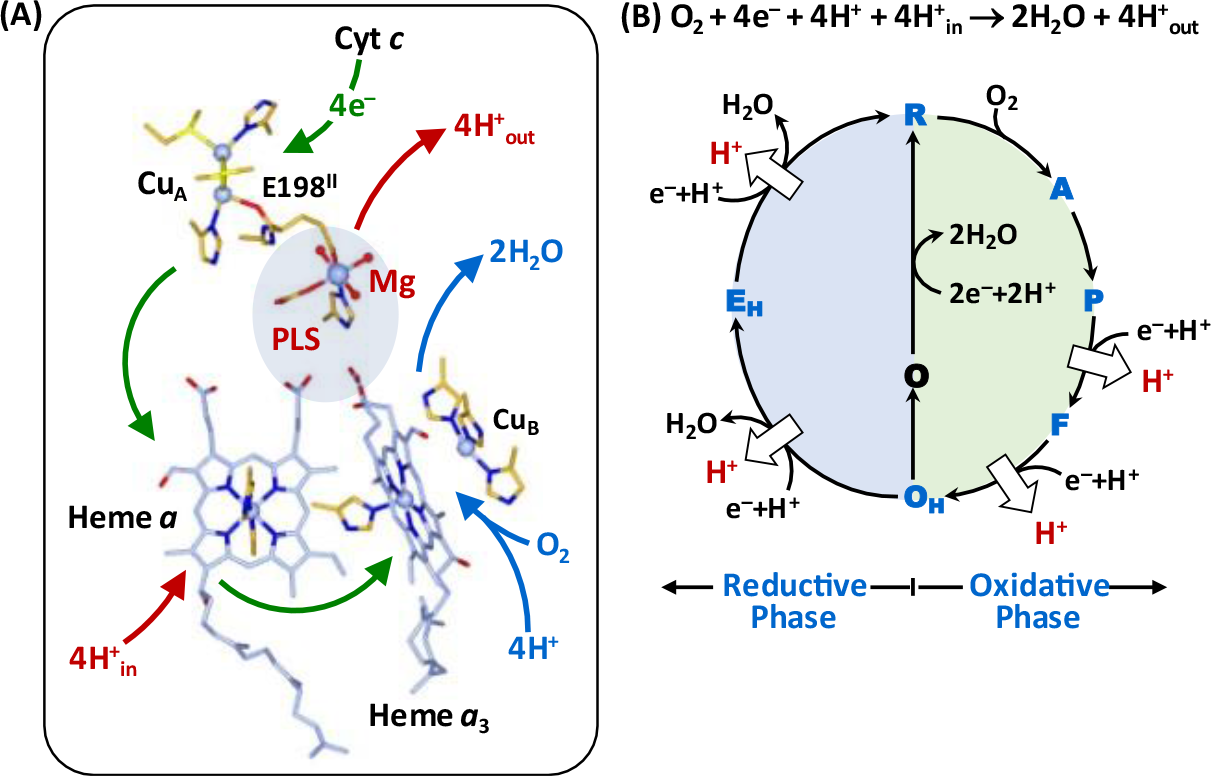
Oxygen reduction reaction catalyzed by bC*c*O. (A) Schematic illustration of the four redox active metal centers in bC*c*O and the electron and proton transfers associated with the O2 reduction reaction. The entry of O2 and four substrate protons into the BNC, as well as the release of the product water molecules out of it, are indicated by the blue arrows. The associated entry of four electrons into the BNC and the translocation of four pumped protons across the membrane are indicated by the green and red arrows, respectively. The putative PLS between heme *a*3 and the Mg center is highlighted by the light blue background. (B) The overall O2 reduction reaction and the associated mechanism. The **P** intermediate is a general term for the **PM** and **PR** intermediates. The entry of the electrons and substrate protons into the BNC and the release of the product water molecules are indicated in each step of the reaction as described in the main text. The coupled proton translocation reactions are indicated by the white arrows. If the **OH** intermediate produced at the end of the oxidative phase is allowed to relax to the resting **O** state, its reduction to **R** does not support proton translocation.

The oxygen reduction reaction catalyzed by C*c*O has been well-characterized^6,16-19^. As illustrated in Fig. 1B, the reaction is initiated by O_2_ binding to the reduced enzyme, **R**, to generate the primary O_2_- complex (**A**). Upon accepting 2 electrons and 2 substrate protons into the BNC, **A** is converted to the oxidized **O**_**H**_ state, *via* the **P** and **F** intermediates. This oxidative phase of the reaction is followed by the reductive phase, where **O**_**H**_ is reduced back to **R** *via* the **E**_**H**_ intermediate, by accepting two additional electrons and 2 substrate protons into the BNC. During the reaction cycle, each time an electron and a substrate proton enter the BNC, a pumped proton is translocated across the mitochondrial membrane, as indicated by the white arrows. Intriguingly, if the metastable **O**_**H**_ state is allowed to relax to the resting **O** state, its reduction to **R**, unlike the **O**_**H**_ →**R** transition, does not drive proton translocation^2,3^, despite the fact that **O** and **O**_**H**_ are redox equivalents.

The structural properties of **O**_**H**_, distinguishing it from the **O** state, have been elusive. Time-resolved resonance Raman spectroscopic studies of bovine C*c*O (bCcO) revealed that the heme *a*_3_ iron in the BNC of the **O**_**H**_ state is coordinated by a hydroxide ion^5,6^, as evidenced by its characteristic ν_Fe-OH_ stretching mode at 450 cm^-1^. In contrast, a comparable ν_Fe-OH_ band has never been identified in the **O** state. As such, it has been thought that the inability of the **O** state to drive proton translocation is at least a partial result of a unique BNC ligation state^16,20,21^. The crystal structure of **O**_**H**_ has not been determined, while those of **O** have been reported for various homologues^22-24^ of C*c*O. It is clear that, in the **O** state, strong electron density associated with exogeneous ligand(s) is present between heme *a*_3_ and Cu_B_ in the BNC; its assignment, however, is controversial. The BNC ligand was first assigned to a peroxide ion bridging the two metal centers^22,25^, but the best fitted O-O bond distance is much longer than that of a typical ferric peroxide species (∼1.48Å). Furthermore, it is uncertain how the peroxide is formed and why it is stable under the equilibrium conditions. Consequently, alternative assignments have been considered. Based on theoretical perspectives, the BNC ligand has been proposed to be a dioxygen^26^ or a water molecule^27^. In contrast, based on structural studies, the ligand in bC*c*O (PDB ID: 7TIE)^28^ and *R. sphaeroides* C*c*O (PDB ID: 2GSM)^23^ has been assigned as a water and a hydroxide coordinated to the heme *a*_3_ and Cu_B_, respectively; although the O-O distance between the two ligands (∼1.9-2.0 Å) is too close for a typical H-bond.

Recently, it was recognized that macromolecular crystallographic structures obtained with intense synchrotron light sources at cryogenic temperatures often suffer from radiation damage problems, in particular for proteins containing redox sensitive metal centers^29-31^. It has been shown that, in the **O** structure of bC*c*O acquired with a typical synchrotron light source, all the four redox active metal centers were reduced, although its polypeptide scaffold remained in a native-like conformation^28,32^. Accordingly, new approaches, such as SFX^4^, have been developed and employed to overcome the radiation damage problems. With SFX, the diffraction patterns of randomly oriented microcrystals suspended in a solution jet are collected sequentially with an X-ray free electron laser (XFEL) before they are destroyed by the intense femtosecond laser pulses. As such, radiation damage-free structures can be obtained at room temperature. Using SFX, Branden and coworkers obtained a radiation damage-free **O** structure of the *ba*_3_ oxidase from *Thermus thermophilus*, based on which it was concluded that the electron density in the BNC was best modeled by a ligand with a single oxygen atom, either a water or a hydroxide ion^24^. Likewise, we have used SFX to determine the radiation damage free **O** structure of bC*c*O^33^; however, we found that it required two oxygen atoms to account for the ligand electron density in the BNC, although the resolution (2.9 Å) was insufficient to clarify the identity of the ligands. On the other hand, Yoshikawa *et al* employed a different point-by-point scanning approach, using an XFEL as a light source, to circumvent radiation damage problems, based on which the BNC ligands of the **O** derivative of bC*c*O were assigned as two co-existing peroxide moieties^34^. Despite these and other efforts, no consensus on the ligand identity has been reached to date. Here, we sought to clarify the structural properties of the **O** derivatives of bC*c*O by using a combination of resonance Raman spectroscopy and SFX.

### Identification of the heme a_3_ ligand by resonance Raman spectroscopy

To determine the identity of the heme *a*_3_ iron ligand, we carried out resonance Raman spectroscopic studies in free soluton. The 413.1 nm output from a krypton ion laser was selected as the excitiation light source to selectively enhance the signals associated with heme *a*_3_. We reasoned that if the heme *a*_3_ iron ligand is a water or a hydroxide ion, it should be able to be exchanged with the solvent water molecules. Accordingly, we incubated the resting **O** derivative of bC*c*O in isotope-substituted H_2^18^_O buffer for 12 hours to ensure complete solvent exchange prior to the resonance Raman measurements. To prevent photoreduction, the laser power was kept low (∼5 mW) and the spectral acquisition time was kept short (3 mins); in addition, the spectra from 6 fresh samples were acquired and summed to improve the signal-to-noise ratio of the spectrum. As a comparison, the spectrum of **O** in naturally abundant H_2_^16^O buffer was obtained in the same fashion.

As shown in Fig. 2, an oxygen isotope sensitive band was identified at 451 cm^-1^ in the H_2_^16^O buffer, which shifted to 428 cm^-1^ in the H_2^18^_O buffer. The isotopic shift of 23 cm^-1^ is consistent with the theoretical shift of a Fe-OH^-^ stretching mode (ν_Fe-OH_) (24 cm^-1^), indicating that the heme *a*_3_ iron ligand of the resting **O** state is a hydroxide ion. It should be noted that, this solvent isotope sensitive mode was not detected in prior studies^6^, possibly due to the complications resulting from laser induced photodamage^35^. The ν_Fe-OH_ mode is identical to that found in the **O**_**H**_ state^5,6^, suggesting that the BNC ligands in **O** and **O**_**H**_ are the same, contrary to the common belief that they are distinct.^16,20,21^ It is noteworthy that this Fe-OH^-^ stretching frequency (451 cm^-1^) is remarkably low as compared to those of other ferric heme species (∼490-550 cm^-1^)^36^, indicating an unusually weak Fe-OH^-^ coordination bond. Hydroxide is generally a strong field ligand for ferric heme iron, which is associated with a strong Fe-OH^-^ bond and a low spin electronic configuration. However, previous resonance Raman^6^ and electron paramagnetic resonance (EPR)^37^ spectroscopic studies revealed that the heme *a*_3_ in the **O** state, like that in the **O**_**H**_ state, has an unusual high spin configuration. We attribute the weak Fe-OH^-^ bond and unique high spin configuration of the heme *a*_3_ to a strong H-bond between the hydroxide ligand and its surrounding environment, as that detected in a hemoglobin from *M. tuberculosis*, which exhibits a similar low Fe-OH^-^ stretching frequency (454 cm^-1^) and a high spin electronic configuration due to a H-bond between the hydroxide ligand and a nearby tyrosine residue^38^. The presence of a strong H-bond to the hydroxide ligand in the BNC of bC*c*O is supported by the structural data discussed below.

**Fig. 2.**
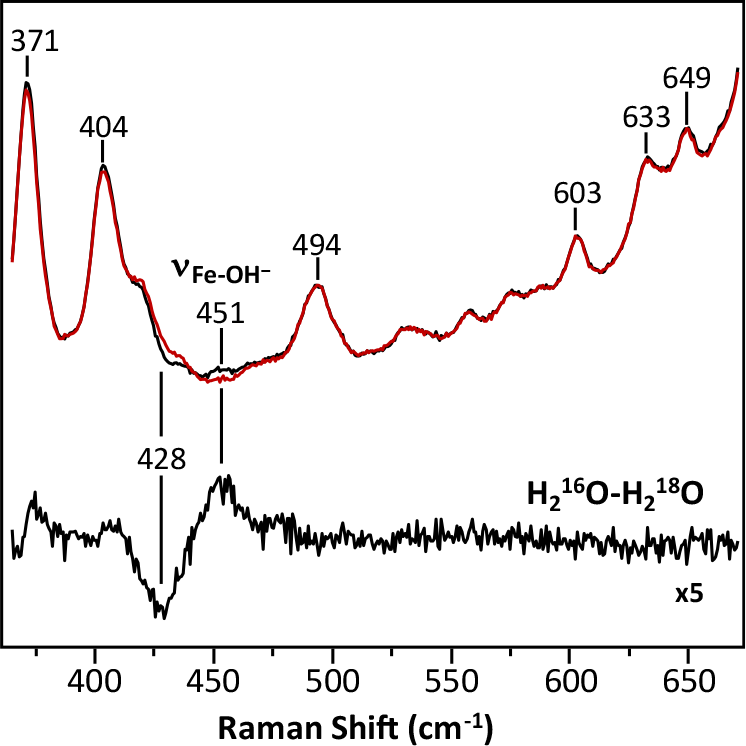
Resonance Raman spectrum of the O state of bC*c*O in H_2_^16^O (black) *versus* H_2_^18^O (red). The H_2_^16^O-H_2_^18^O difference spectrum (expanded by 5 fold) is shown at the bottom. The oxygen sensitive mode in the H_2_^16^O sample centered at 451 cm^-1^ that shifted to 428 cm^-1^ in the H_2_^18^O sample is assigned to the νFe-OH mode.

### Determination of the structure of the O state by SFX

For the SFX measurements, microcrystals of bC*c*O were prepared in the resting **O** state using a previously reported protocol^33^. To ensure the homogeneity of the **O** state, the microcrystals were reduced under anaerobic conditions and then allowed to turn over by exposing to O_2_ and subsequently relax back to the **O** state by incubating overnight in a stabilizing solution. The suspension of the microcrystals was loaded into a gas-tight syringe and injected into the XFEL beam as a free solution jet with a single capillary MESH injector^39,40^. The XFEL beam, perpendicular to the solution jet, was directed into the tip of the Taylor cone formed at the output of the MESH injector. X-ray diffraction was collected for approximately 2 hours, from which 84,736 indexable diffraction patterns were selected for structural analysis. The structure was solved and refined to a resolution of 2.38 Å (Extended Data, Table 1). The structural markers for the oxidation states of the four redox centers confirm that the enzyme is in the fully oxidized **O** state (Extended Data Fig. 2).

### Structural characterization of the BNC

In the F_O_-F_C_ electron density map associated with the SFX data (Fig. 3A), a large 2-lobe electron density is evident in the BNC, indicating the presence of ligands coordinated to heme *a*_3_ iron and Cu_B_. As guided by the resonance Raman data (Fig. 2), we modeled the heme *a*_3_ ligand density with a OH^-^ ion (Fig. 3B). In addition, we modeled the Cu_B_ ligand density with a water molecule, as the coexistence of two negatively charged hydroxide ions in the BNC is expected to be energetically unfavorable. The occupancy of the ligands is confirmed by the polder maps shown in Fig. 3C. The Fe-OH^-^, Cu_B_-H_2_O and Fe-Cu_B_ bond lengths, which were unrestrained during the refinement, are determined to be 1.90, 2.14 and 4.74 Å, respectively. The O-O distance between the two oxygen ligands is 2.53Å, which is shorter than a typical H-bond, but is consistent with a strong low-barrier H-bond^41,42^. This strong H-bond between the two ligands plausibly weakens the ligand field strength of the hydroxide and destabilizes the Fe-OH^-^ bond, thereby accounting for the high spin electronic configuration and the low Fe-OH^-^ stretching frequency revealed by the spectroscopic studies. Taken together our data demonstrate for the first time that the heme *a*_3_ iron and Cu_B_ in the **O** state, like those in the **O**_**H**_ state, are coordinated by a hydroxide ion and a water molecule, respectively.

**Fig. 3.**
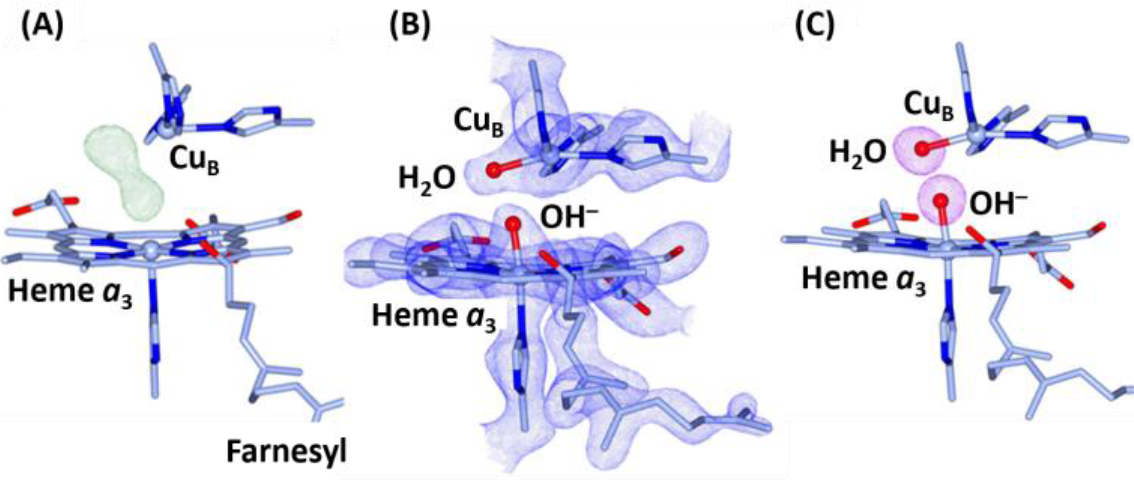
Electron density maps of the BNC in the O state of bC*c*O. (A) The FO-FC electron density map (contoured at 7.0 σ) showing clear 2-lobe electron density associated with the BNC ligands. (B) The 2FO-FC electron density map (contoured at 2.5 σ) obtained with the electron density modeled with a hydroxide ion coordinated to the heme *a*3 iron and a water molecule coordinated to CuB. (C) The Polder map (contoured at 7.0 σ) associated with the BNC ligands.

### Protonation state of Y244

Cu_B_ in the BNC is coordinated by three histidine ligands, one of which (H240) forms a covalent bond with Y244 by a posttranslational modification (Fig. 4). It is well-established that, during the catalytic reaction, Y244 temporarily donates an electron and a proton to the heme *a*_3_ bound O_2_ to promote the O-O bond scission, by forming a tyrosyl radical^43^, and that its re-reduction to a tyrosinate and re-protonation back to its neutral protonated form are tightly coupled to the ensuing electron and proton transfer processes^20,43,44^.

**Fig. 4.**
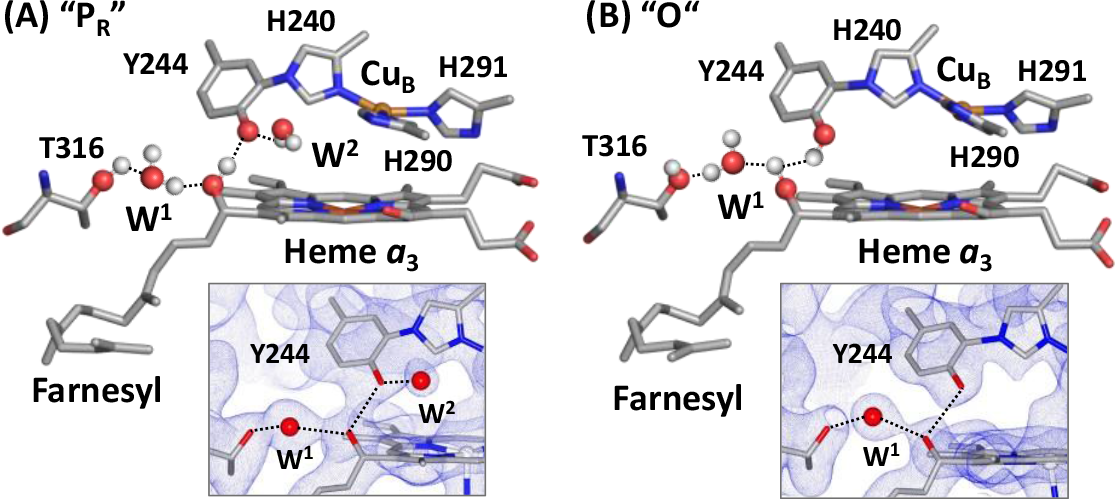
Protonation state of Y244 in the PR (A) and O (B) states. The post translationally modified Y244 forms a H-bonding network with the OH group of the farnesyl side chain of heme *a*3, a water molecule (W^1^) and T316. In the **PR** state, an additional water, W^2^, is recruited into the BNC to stabilize the tyrosinate configuration of Y244. This water is absent in the **O** state reported here, as evident in the 2FO-FC electron density map (contoured at 1.0 σ) shown in the lower inset, signifying that Y244 is in the neutral protonated state. For clarity the BNC ligands are not shown. The oxygen atom and hydrogen atoms are shown as red and white spheres.

Previously, with time-resolved SFX, we showed that, in the **P**_**R**_ intermediate, Y244 is in the tyrosinate form^33^. Next to the Y244, there is a water molecule within a H-bonding distance from the oxygen atom of the tyrosinate (indicated as W^2^ in Fig. 4A), which is not present in the reduced **R** state (PDB IDs: 6NMF and 7THU) or the CO complex (PDB ID: 5W97, a structural analog of the **A** state), where Y244 is in the neutral protonated form. It suggests that, in the **P**_**R**_ state, W^2^ is recruited into the BNC to stabilize the tyrosinate configuration of Y244, similar to the water rearrangement induced by tyrosine deprotonation found in a photosynthetic reaction center from *B. viridis* _45_. Intriguingly, our current work reveals that W^2^ is not present in the **O** state (Fig. 4B), signifying that Y244 is in the neutral protonated form. Although the structures of the **F** and **O**_**H**_ intermediates have not been determined, infrared spectroscopic studies demonstrate that Y244 in both **F** and **O**_**H**_, like that in the **P**_**R**_ intermediate, is deprotonated^44^, indicating that during the **P**_**R**_→**F**→**O**_**H**_ transition, Y244 remains in the tyrosinate configuration. This scenario is in good agreement with recent structural data showing the presence of W^2^ in the mixed oxygen intermediates of bC*c*O^21,46^. Our current data further demonstrate that the **O**_**H**_→**O** transition is associated with the protonation of the tyrosinate (Y244).

### The oxygen reaction cycle

With the clarification of the **O** structure, we propose a complete reaction cycle of C*c*O as illustrated in Fig. 5. The reduced **R** state first binds O_2_ to form the primary oxy intermediate **A**, with a Fe^3+^-O^2-^ electronic configuration^47^. By accepting an electron and a proton from Y244, the O-O bond in **A** is heterolytically cleaved, leading to the formation of the putative **P**_**M**_ intermediate, where one oxygen remains on the heme *a*_3_ iron, in a ferryl (Fe^4+^=O^2-^) configuration, and the other oxygen is coordinated to Cu_B_ as a hydroxide, while Y244 is converted to a neutral tyrosyl radical. The entry of one electron into the BNC leads to the conversion of **P**_**M**_ to **P**_**R**_, where the tyrosyl radical is reduced to a tyrosinate. The **P**_**M**_→**P**_**R**_ transition is associated with the entry of a new water (W^2^, not shown in Fig. 5 for clarity), which stabilizes the tyrosinate configuration of Y244^33^. The subsequent entry of a substrate proton into the BNC further transforms **P**_**R**_ to **F**, where the hydroxide ligand of Cu_B_ is protonated to a water, which strengthens the iron-oxygen bond as evidenced by the shift of the frequency of the Fe^4+^=O^2-^ stretching mode from 785 to 804 cm^-1^.^48^ Finally, the entry of an additional electron and a substrate proton into the BNC converts **F** to **O**_**H**_, where the Fe^4+^=O^2-^ moiety is reduced to the Fe^3+^-OH^-^ species^5,6,48^. During this oxidative phase, one pumped proton is translocated during each of the **P**_**R**_→**F** and **F**→**O**_**H**_ transitions (as indicated by the white arrows) and, throughout the **P**_**R**_→**F**→**O**_**H**_ transformation, Y244 remains in the tyrosinate form.

**Fig. 5.**
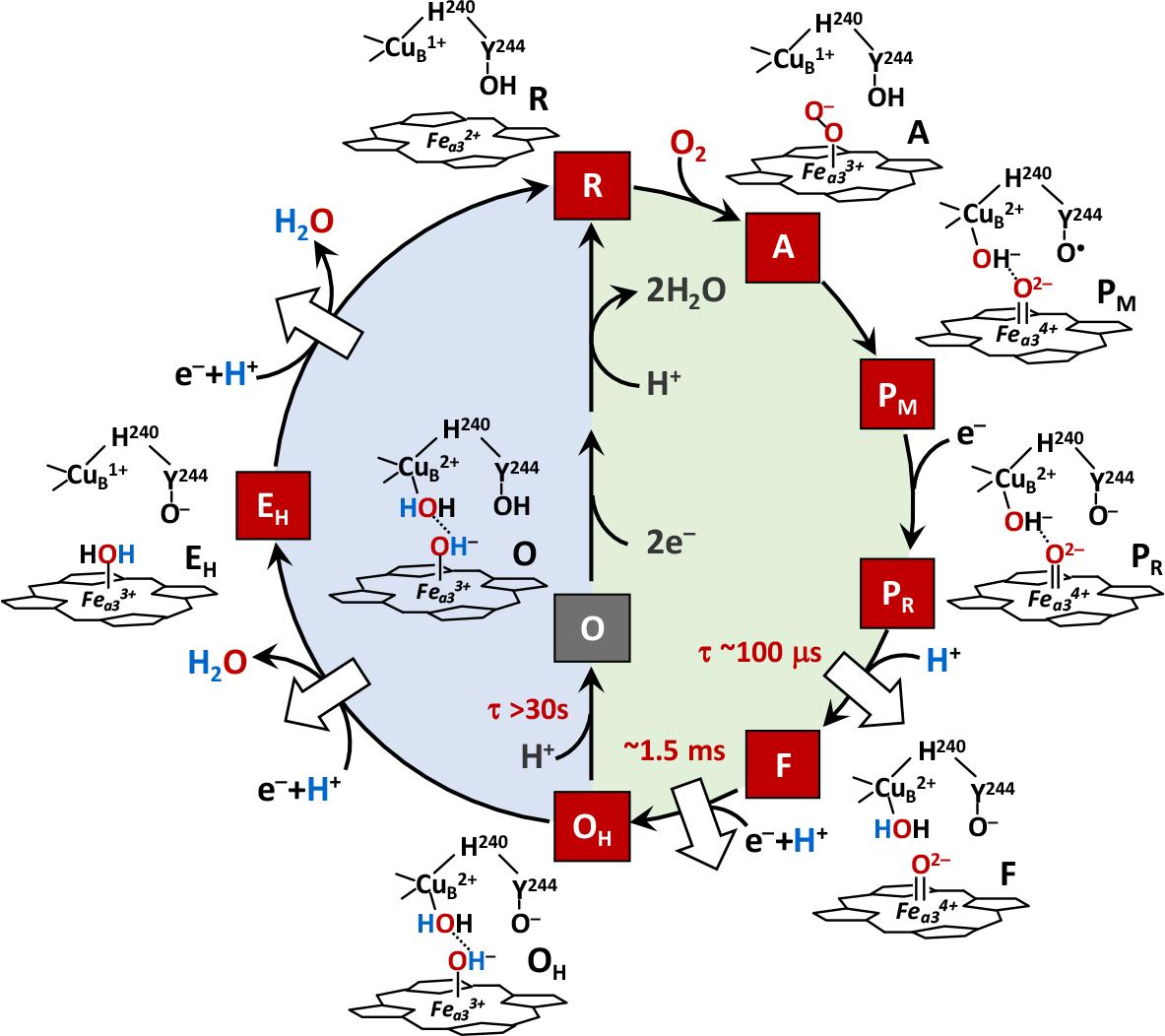
Hypothesized ligand transformation in the BNC during the C*c*O reaction cycle. The oxidative and reductive phase of the reaction cycle are highlighted with green and blue backgrounds, respectively. The white arrows indicate the proton translocation reactions associated with the reaction cycle. The lifetimes associated with the **PR**→**F, F**→**OH** and **O**H→**O** transitions are indicated in red.

In the ensuing reductive phase, an additional electron and a substrate proton enter the BNC, leading to the conversion of **O**_**H**_ to **E·**_**H**_, where Cu_B_ is reduced from the cupric to cuprous state and the hydroxide ligand of the heme *a*_3_ is protonated to a water. At the same time, the water ligand of Cu_B_ is released out of the BNC. The entry of another electron and a substrate proton into the BNC leads to the conversion of **E**_**H**_ to **R**, where the heme *a*_3_ iron is reduced from the ferric to ferrous state and its water ligand is released out of the BNC; at the same time the tyrosinate is protonated back to its neutral form. During this reductive phase, one additional pumped proton is translocated during each of the **O**_**H**_→**E**_**H**_ and **E**_**H**_→**R** transitions. Under electron deficient conditions, the **O**_**H**_ intermediate produced at the end of the oxidative phase can spontaneously and slowly relax to the resting **O** state^3^. Our current data revealed that during the **O**_**H**_→**O** transition, the tyrosinate (Y244) is protonated to the neutral form, leading to the dissociation of the W^2^ that stabilizes the tyrosinate, while the BNC ligands remain unchanged.

### Implications on proton translocation

It has been shown that the entry of the first two substrate protons into the BNC, associated with the **P**_**R**_→**F** and **F**→**O**_**H**_ transitions, is mediated by the D-channel and occurs rapidly (within ∼100 μs and 1.5 ms, respectively) ^1^. Similarly, the entry of the other two substrate protons into the BNC, associated with the **O**_**H**_→**E**_**H**_ and **E**_**H**_→**R** transitions, also takes place quickly (on the submillisecond time scale), although it is mediated by a different channel, the K-channel^49^. In sharp contrast, the off-pathway **O**_**H**_→**O** transition is sluggish and does not reach completion until ∼30 s,^3^ suggesting that the tyrosinate (Y244) is likely protonated by an adventitious proton from its surrounding environment, rather than a substrate proton from the K-channel.

Based on the electroneutrality principle proposed by Rich *et al*^50,51^, during the catalytic reaction, the entry of each electron into the BNC (see the green arrows in Fig. 6A) is charge-compensated by the entry of one substrate proton from the D or K-channel (blue arrows). The redox energy thereby derived is then used to drive the release of a pumped proton into the P-side of the membrane (red arrows) from the PLS, which is preloaded with the pumped proton(s) *via* the D or H-channel. As illustrated in Fig. 5, Y244 in all the intermediate states active in proton translocation (**P** and **E**_**H**_) is in the deprotonated tyrosinate form, suggesting that the premature protonation of the tyrosinate (Y244) during the **O**_**H**_→**O** transition, without the input of any electron into the BNC, perturbs the charge balance in the BNC of the **O** state, thereby disabling its ability to release pumped protons out of the PLS upon its reduction (Fig. 6B). This scenario aligns well with recent computational studies demonstrating the importance of deprotonated forms of Y244 in modulating the energy landscape of the C*c*O reaction to promote the coupling of the oxygen reduction chemistry to proton translocation^52^.

**Fig. 6.**
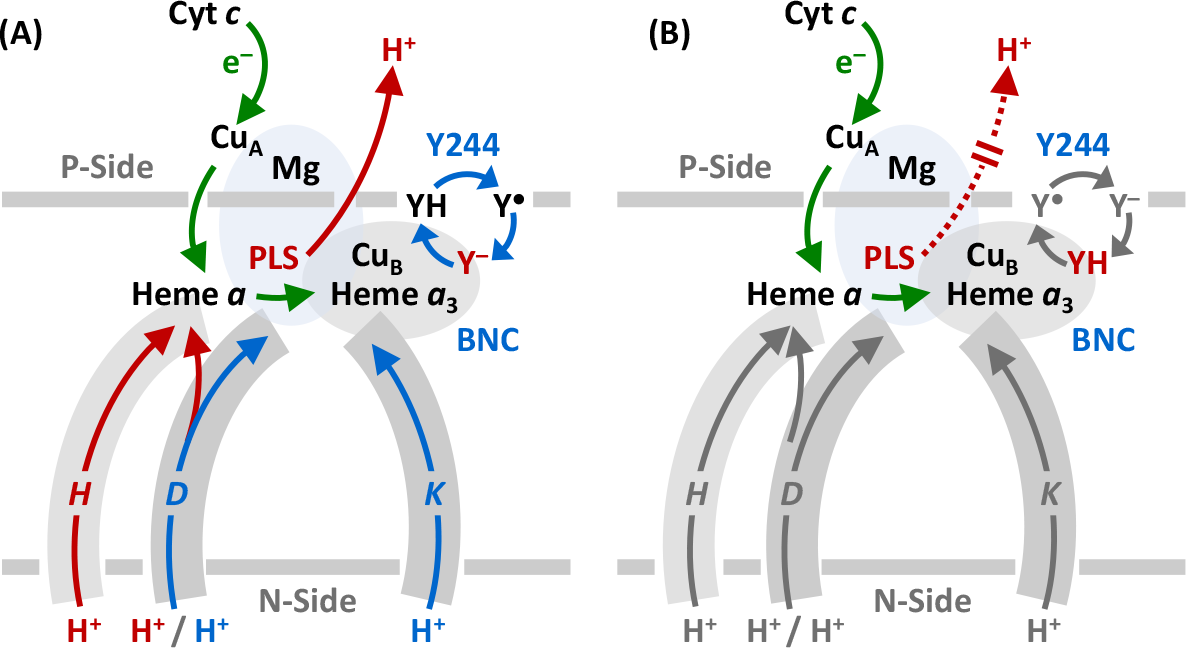
Functional role of Y244 during the C*c*O reaction cycle. (A) During the active turnover, Y244 cycles between the neutral form (YH), tyrosyl radical form (Y^●^) and deprotonated tyrosinate form (Y^**–**^) as depicted by the blue cycle. In all the intermediate states active in proton translocation, Y244 is in the Y^**–**^ form (in red), which ensures the tight coupling of electron transfer (green arrows) and substrate proton transfer (blue arrows), *via* the D and K-channel into the BNC, to drive the proton translocation (red arrows) *via* the D or H channel from the N to P-side of the membrane. (B) The protonation of the tyrosinate to the YH form (red) in the **O** state disables the proton translocation upon its reduction to **R**.

### Conclusions

Previous studies reveal that **O** and **O**_**H**_ are redox equivalents, but only the latter, not the former, is capable of translocating protons upon reduction. With a combination of resonance Raman spectroscopy and SFX, we now clarify that the heme *a*_3_ iron and Cu_B_ in the BNC of the **O** state, like those in the **O**_**H**_ state, are coordinated by a hydroxide ion and water, respectively; in addition, we show that the post-translationally modified Y244 is in the neutral protonated form, which distinguishes the **O** state from the **O**_**H**_ state, where Y244 is in the deprotonated tyrosinate form. Our data underscore the pivotal role of the deprotonated form of Y244, a residue fully conserved in the CcO family of enzymes, in energizing the enzyme for proton translocation, as supported by reported computational studies^52^.

## Materials and Methods

### Protein sample preparation

Bovine C*c*O was isolated from bovine hearts by a standard procedure^53,54^. In the final step of the purification, the enzyme was crystallized by concentration on an Amicon concentrator equipped with a 200 kDa molecular cutoff filter. The crude crystals were harvested and redissolved in 40 mM pH 6.8 phosphate buffer containing 0.2% decylmaltoside to generate the protein stock for the resonance Raman measurements and microcrystal preparation for the SFX studies.

### Resonance Raman measurements

To prepare the sample in H_2^18^_O, a concentrated bC*c*O sample (in 40 mM pH 7.4 phosphate buffer containing 0.2% decylmaltoside) was diluted by 10-fold with H_2^18^_O containing the same buffer and then incubated overnight prior to the resonance Raman measurements. The final protein concentration was 30 µM. An equivalent sample in H_2_^16^O was prepared in the same fashion as a comparison. Resonance Raman spectra were obtained by using 413.1 nm excitation from a Kr ion laser (Spectra Physics, Mountain View, CA). The laser beam was focused to a ∼30 µm spot on the spinning quartz cell rotating at ∼1000 rpm. The scattered light, collected at right angle to the incident laser beam, was focused on the 100 µm wide entrance slit of a 1.25 m Spex spectrometer equipped with a 1200 grooves/mm grating (Horiba Jobin Yvon, Edison, NJ), where it was dispersed and then detected by a liquid-nitrogen cooled CCD detector (Princeton Instruments, Trenton, NJ). A holographic notch filter (Kaiser, Ann Arbor, MI) was used to remove the laser scattering. The Raman shift was calibrated by using indene (Sigma).

It has been shown that high laser power can induce photoreduction linked artifacts in the resonance Raman spectra of bC*c*O^35^, which prevented identification of the Fe-OH stretching mode in a prior study^6^. As such, we kept the laser power low (∼5 mW at the sample) and limited the laser exposure time to 3 minutes for each sample to avoid photoreduction and photodamage. A total of 6 spectra were acquired from fresh samples and averaged to improve the signal to noise ratio of the spectrum. To assure that artifacts were not present, spectra of the H_2_^16^O and H_2^18^_O buffers alone were obtained. It was confirmed that no isotopic differences in the 400 – 450 cm^-1^ window were detected.

### Microcrystal preparation

The microcrystals were prepared with a previously reported method^33^. The crystal growth was initiated by mixing the protein stock with the precipitant solution (0.2% decylmaltoside and 2.5% PEG4000 in 40 mM pH 6.8 phosphate buffer) and a seeding solution (prepared by crushing and sonicating large crystals in the mother solution). The microcrystals were allowed to grow at 4° C for ∼36 hours before they were harvested and characterized by polarized optical microscopy. The microcrystals have a planar shape with approximate dimensions of ∼20 × 20 × 4 μm. To obtain a homogeneous sample of the oxidized **O** crystals for the SFX measurements, the microcrystals were reduced anaerobically by a minimum amount of dithionite in a glove box and then thoroughly washed with the mother solution to remove excess dithionite and its oxidized products. The reduced microcrystals were then exposed to O_2_ to initiate the enzyme turnover. They were allowed to relax in a stabilizing solution (0.2 % decylmaltoside and 6.25% PEG-4000 in 40 mM pH 6.8 sodium phosphate solution) at 4 °C for ∼24 hours prior to the SFX measurements.

### SFX data acquisition and structural analysis

The SFX measurements were conducted at the Macromolecular Femtosecond Crystallography (MFX) end station of the Linac Coherent Light Source (LCLS) at the SLAC National Accelerator Laboratory. A bC*c*O microcrystal slurry was loaded into a gas tight syringe and driven by a HPLC pump into a 100 µm diameter capillary of a Microfluidic Electrokinetic Sample Holder (MESH) injector,^39,40^ which has recently been developed as a useful and reliable technique for delivering microcrystal slurries into the XFEL beam^55-59^. A high voltage (∼+2500 V) was applied to the sample at the entrance of the capillary against a grounded waste collector. It was used to electro-focus the microcrystal jet down to a Taylor cone at the capillary output. The 10 keV / 30 fs pulses from the X-ray free electron laser (XFEL) intersected the microcrystal slurry at the tip of the Taylor cone prior to its development into a thin jet, as the jet was too unstable for data collection, while the thick region of the cone gave too many multiple hits and a high background. The diameter of the XFEL beam was ∼3 μm. The sample flow rate was set to ∼3 µl/min. A series of diffraction patterns from randomly orientated microcrystals were collected with a Rayonix MX340-XFEL CCD detector at a 30 Hz rep rate.

The quality and hit rate of the SFX data were monitored in real time using OM^60^. The data were collected for approximately 2 hours. *Psocake*^61,62^ was used to determine the initial diffraction geometry and find crystal hits. *CrystFEL*’s indexing program was used to index the crystal hits^63,64^. A total of 84,736 crystals were indexed. The initial structure was solved with molecular replacement with Phaser-MR through the CCP4 program suite^65^ using a 1.9 Å resolution structure of bC*c*O (PDB ID: 7TIE) as the search model. Waters and BNC ligands were excluded from the search model. Further model building was performed using Coot^66^. Structure refinements were done using Refmac5 and PDB-Redo^67^. The final structure was refined to a resolution of 2.38 Å (see Extended Data Table 1) (PDB ID: 8GCQ).

## ACKNOWLEDGEMENTS

The SFX experiments were carried out at the Linac Coherent Light Source (LCLS) at the SLAC National Accelerator Laboratory. LCLS is an Office of Science User Facility operated for the US Department of Energy Office of Science by Stanford University. Use of the LCLS, SLAC National Accelerator Laboratory, is supported by the U.S. Department of Energy, Office of Science and Office of Basic Energy Sciences under Contract No. DE-AC02-76SF00515. The HERA system for in helium experiments at MFX was developed by Bruce Doak and funded by the Max-Planck Institute for Medical Research. This work was supported by National Science Foundation (NSF) CHE-1404929 (D.L.R. and S.-R.Y.) and National Institutes of Health (NIH) grants S10 OD023453, P41 GM139687, GM126297 (D.L.R. and S.-R.Y.) and GM115773 (S.-R.Y.).

## Author Contributions

SRY, II and DLR designed experiments; II isolated and crystallized bCcO and measured and interpreted the resonance Raman spectra; RGS designed and operated the MESH injector; RES prepared the injector auxiliary equipment; FP, SL, BH, FRM and SB collected the SFX data; ZS, AP, CW and CHY processed the SFX data; SRY, II, CHY and DLR analyzed and interpreted the SFX data; SRY and DLR drafted the manuscript and all authors contributed to the revisions.

