## Supplemental Data for "Structural basis for functional properties of cytochrome *c* oxidase"

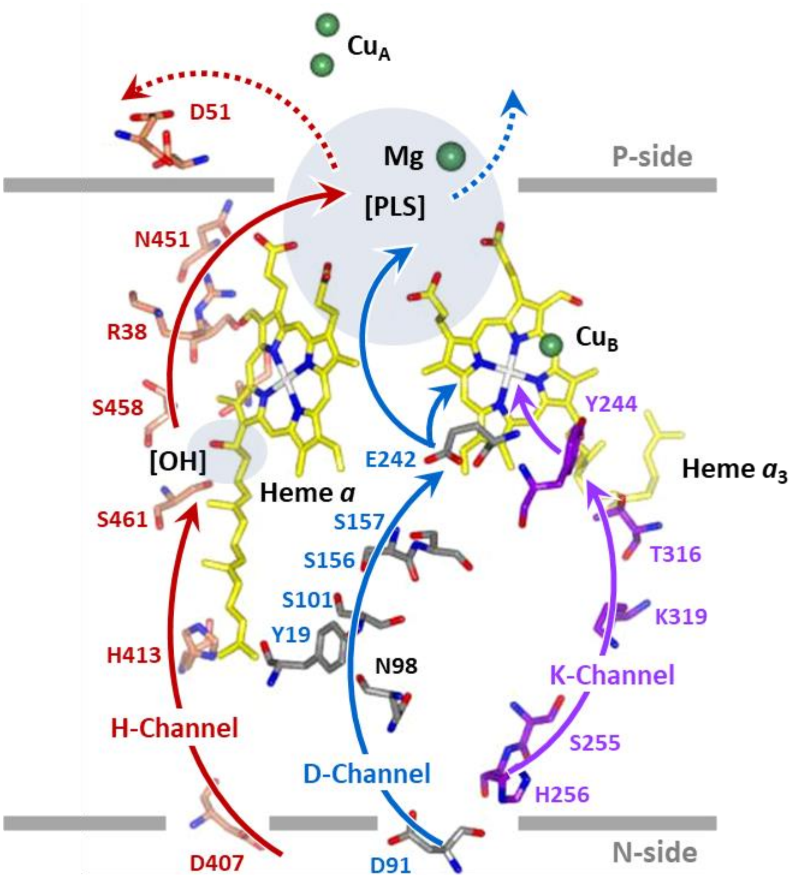


**Extended Data Fig. 1. Proton transfer channels in bCcO.** Based on structural and mutagenesis studies of bacterial enzymes, two proton transfer channels, the D and K-channels, have been proposed in C*c*O[1]. The D-channel, starting at D91 (bC*c*O numbering) and ending at E242 (which serves as a diverter valve), delivers either substrate protons to the BNC during the oxidative phase of the catalytic cycle, or pumped protons to the PLS for subsequent translocation to the P-side of the membrane. The K-channel, starting at H256 and ending at Y244, delivers substrate protons to the BNC during the reductive phase of the catalytic cycle. An additional channel, the H-channel, starting at D407 and ending at D51, was proposed in bC*c*O, which translocates pumped protons to the P-side of the membrane *via* the PLS, as gated by either a water pool[2] or the hydroxide group of the farnesyl sidechain (indicated as [OH])[3] of heme *a*. The role of the H-channel, however, remains controversial as it is not conserved in bacterial CcOs[4,5].

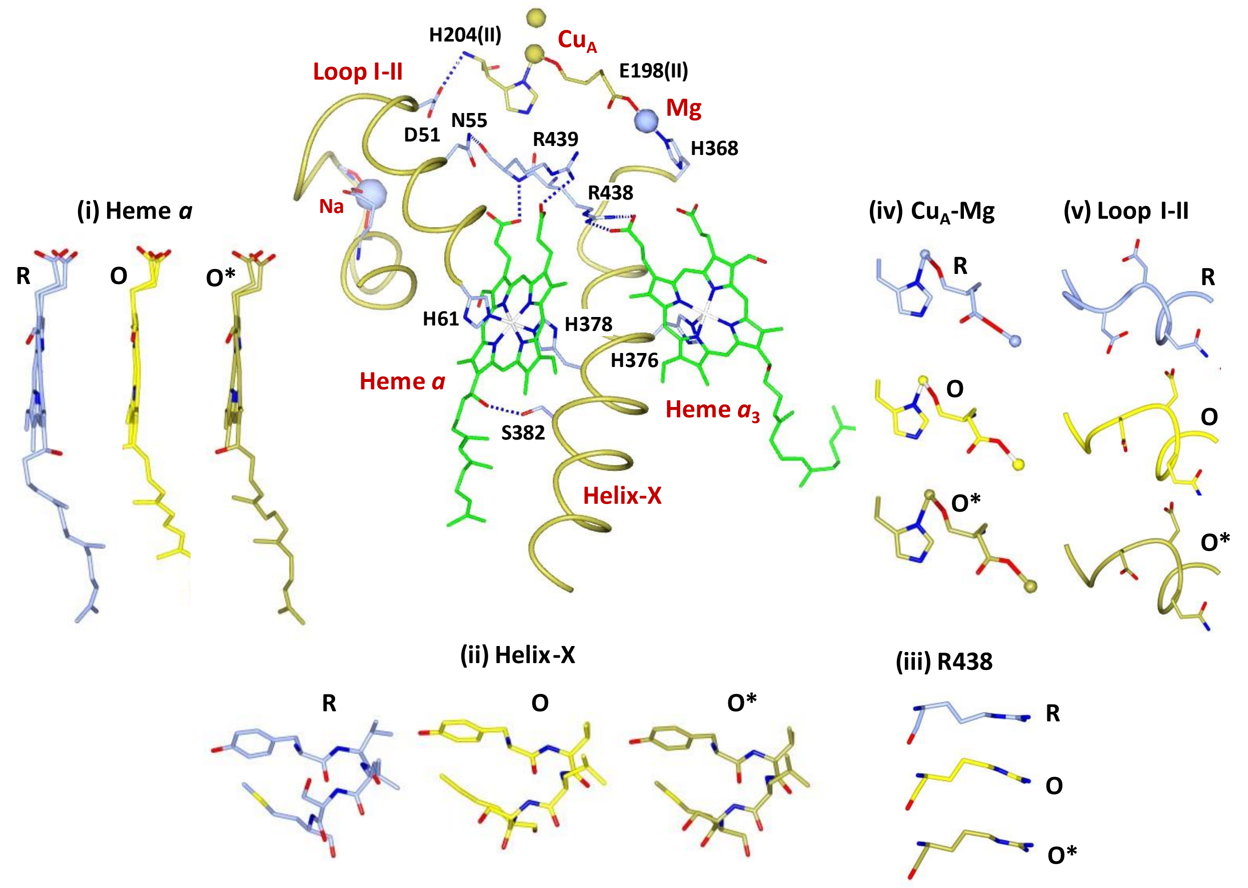


**Extended Data Fig. 2. SFX structure of the O derivative of bC*c*O in the redox sensitive regions.** Inset (i)-(v) show the structural comparison of **O** with the oxidized (**O***) and reduced (**R**) enzyme in the redox sensitive regions, which demonstrate that the structure of **O** is in the fully oxidized conformation. The PDB IDs of the **O*** and **R** are 7TIE and 7THU, respectively.

**Extended Data Table 1**. Crystallographic data collection and refinement statistics. The values in the parentheses are for the outer shell.

| **Data Collection** | |
|---|---|
| Collection Temperature | 293 K |
| Space Group | $P2_12_12_1$ |
| Dimensions: a, b, c (Å) | 178.6, 189.5, 211.1 |
| Dimensions α, β, γ (°) | 90, 90, 90 |
| Resolution (Å) | 33.0 - 2.38 |
| I/σI | 4.59 (1.78) |
| CC* | 0.999 (0.300) |
| Redundancy | 2742 (2286) |
| Completeness (%) | 100 (100) |
| Number of Indexed Hits | 84,736 |
| **Refinement** | |
| Resolution (Å) | 35.00 – 2.38 |
| Unique reflections | 270, 854 (19,803) |
| $R_{work}/R_{free}$ | 0.22/0.26 (0.28/0.31) |
| Number of atoms | 31,465 |
| Average B factor (Å$^2$) | 33.0 |
| **R.m.s. deviations** | |
| Bond lengths (Å) | 0.0114 |
| Bond angles (°) | 1.742 |
| **Ramachandran statistics (%)** | |
| Favored regions | 89.24 |
| Outliers | 1.03 |
| Molprobity score | 2.86 |
